# Neratinib Synergizes with Trastuzumab Antibody Drug Conjugate or with Vinorelbine to Treat HER2 Mutated Breast Cancer Patient Derived Xenografts and Organoids

**DOI:** 10.1101/2023.12.19.572069

**Authors:** Shunqiang Li, Tina M. Primeau, Maureen K. Highkin, Stephanie L. Pratt, Ashley R. Tipton, Nagalaxmi Vemalapally, John Monsey, Yu Tao, Jingqin Luo, Ian S. Hagemann, Chieh-Yu Lin, Lisa D. Eli, Cynthia X. Ma, Ron Bose

## Abstract

HER2 (*ERBB2*) is a major therapeutic drug target in breast cancer and The Cancer Genome Atlas (TCGA) Breast Cancer project and other studies have identified HER2 activating mutations in breast cancers without HER2 gene amplification. HER2 activating mutations occur in 2-5% of metastatic breast cancer patients (MBC), and clinical trials have shown that the irreversible pan-HER tyrosine kinase inhibitor, neratinib, produces a 31-40% clinical benefit rate for HER2 mutated MBC patients. We developed breast cancer patient-derived xenografts (PDX) from ER+, HER2 mutated MBC patients and used them to test neratinib-based drug combinations. Using organoid culture of these PDX breast cancer cells, we performed rapid, high-throughput *ex vivo* screening assays to test novel drug combinations. These organoid culture experiments identified drug synergy with the neratinib plus ado-trastuzumab emtansine (T-DM1) and neratinib plus vinorelbine combinations and we validated these results with *in vivo* PDX experiments.

**Statement of Significance:** PDX’s are a ready source of human cancer organoids, and with thousands of PDX’s already available worldwide, PDX derived organoids (PDxO’s) can dramatically accelerate cancer drug testing. This strategy of PDxO drug testing is particularly useful for rare cancer subtypes or mutations to identify the most promising treatment strategies for clinical trials testing.

## Introduction

HER2 (*ERBB2*) is a major therapeutic drug target in breast cancer. Trastuzumab and other HER2 targeted drugs have dramatically improved outcomes for patients with HER2 gene amplified breast cancers (1,2). One of the major findings of The Cancer Genome Atlas (TCGA) Breast Cancer project was the identification of HER2 somatic mutations in breast cancers without HER2 gene amplification (3,4). A subset of HER2 mutations were shown to be activating mutations that occur in 2-5% of patients with metastatic breast cancer (MBC) (4,5). The most common HER2 activating mutations in breast cancer include missense mutations or small in-frame insertions in the kinase domain or missense mutations at codons 309-310 of the extracellular domain (4,6,7). In preclinical studies, these HER2 activating mutations confer resistance to hormonal therapy or the first generation EGFR/HER2 inhibitor lapatinib, but can be potently inhibited with the irreversible, pan-HER tyrosine kinase inhibitor, neratinib (4,5,8). Two prospective clinical trials, the MutHER phase II clinical trial for HER2 mutations in MBC and the SUMMIT basket trial of neratinib for HER family mutations in cancer, have both demonstrated that neratinib monotherapy in HER2 mutated MBC patients produces a 31-40% clinical benefit rate (6,7). While this is strong evidence for single agent activity of neratinib for this molecular subset of MBC, the progression free survival was modest and both trials added fulvestrant to neratinib in the hormone receptor positive (ER+) population because of crosstalk between HER2 and ER signaling, and to increase clinical efficacy (9,10). However, the relative low frequency of these mutations has slowed progress and more rapid means to accelerate drug combination testing on HER2 mutated MBC is needed.

Cancer organoids are suitable for high-throughput drug screening and offer a solution to this problem. Organoids are a novel, 3D *ex vivo* culture system to grow adult stem cells and primary human cancers (11,12). Organoid culture is based on the three dimensional (3D) culture systems developed by Bissell, Werb, and others (13–15). In 2009, Clevers and colleagues reported organoid culture conditions for adult, intestinal epithelial stem cells (16) and in 2018, they reported culture conditions for growing primary human breast cancer samples *ex vivo* (17). While cancer organoids made directly from patient samples offer great promise for the future, another ready source of human cancer organoids are existing PDX’s. Over 500 breast cancer PDX’s exist worldwide (18,19) and across all cancers types, the numbers of existing PDX’s likely numbers in the thousands (20,21).

In this study, we established breast cancer organoids from two PDX’s from HER2 mutated, ER positive MBC patients. Using these organoids, neratinib drug combinations were rapidly tested in 96 and 384 well plate experiments. These results were then validated with *in vivo* PDX experiments. These results identify two synergistic, drug combination regimens that can be considered for future clinical trials.

## Results

### Establishing HER2 Mutated Breast Cancer PDX’s and Testing Neratinib plus Fulvestrant

We established patient derived xenografts (PDX) from two patients with HER2 mutated, ER positive and HER2 non-amplified MBC. These samples are designated WHIM51 and WHIM64 (WHIM=<u>W</u>ashington University <u>H</u>uman In <u>M</u>ouse). The prior hormonal therapies and chemotherapies that these two patients received are listed in table 1. WHIM51 is an invasive ductal breast carcinoma that has a HER2 exon 20 insertion mutation at amino acid 776 (*ERBB2* A775_G776insYVMA), which is located in the HER2 tyrosine kinase domain. In comparison, WHIM64 is an invasive lobular breast carcinoma that has a HER2 L869R missense mutation in the tyrosine kinase domain. Both of these HER2 mutations have been previously characterized and are known activating mutations (4,22,23).

**Table 1:**
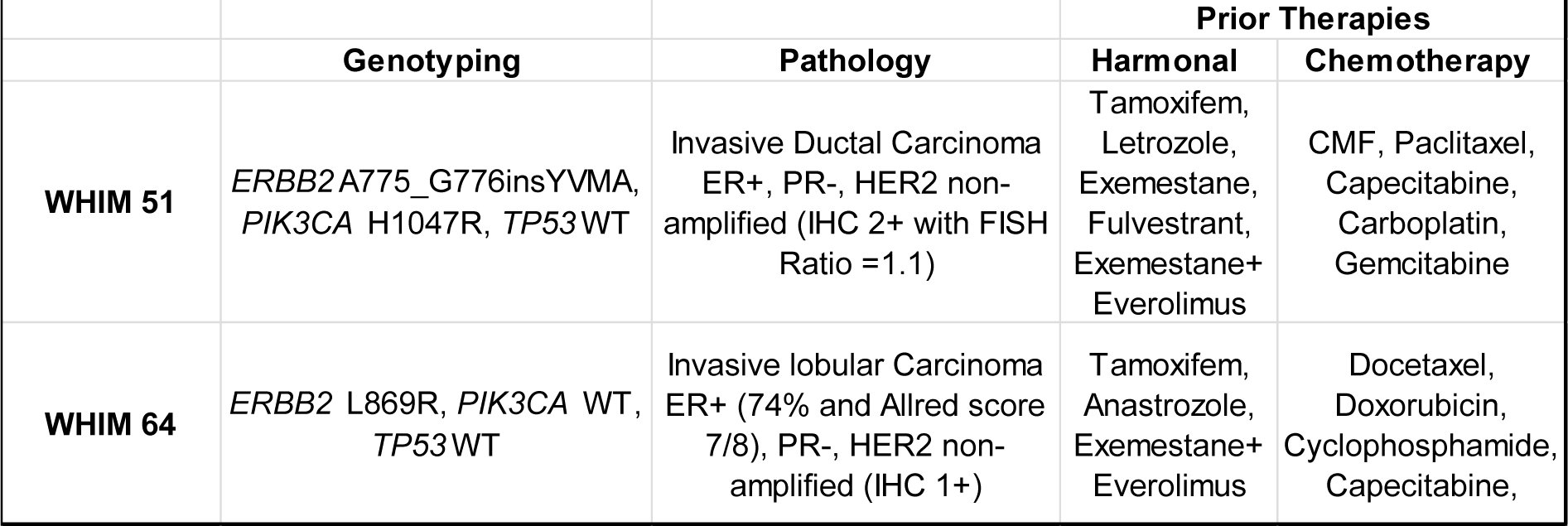
WHIM 51 and WHIM 64 information. Genotype, pathology and treatment history of HER2 mutated breast cancer PDX’s used in this study. CMF = Cyclophosphamide, methotrexate,5-FU.

The MutHER and SUMMIT clinical trials tested the combination of neratinib plus fulvestrant for HER2 mutated, ER positive metastatic breast cancer (9,10). Therefore, we tested this combination in both WHIM51 and WHIM64 (figure 1A-B). Both PDX’s were very sensitive to single agent neratinib. Single agent fulvestrant caused tumor regression of WHIM51 whereas it had a largely static effect on the tumor size of WHIM64. In both of these experiments, there was no statistical difference between the neratinib plus fulvestrant combination versus neratinib monotherapy (p=.99 and p=.69 for WHIM51 and WHIM64, respectively). Because the WHIM64 tumors became undetectable by day 10 when treated with neratinib 40 mg/kg, we performed a dose titration experiment of neratinib (figure 1C) in order to select a dose that would facilitate testing of neratinib drug combinations. Based on this experiment, we selected a dose of neratinib 20 mg/kg for subsequent experiments with WHIM64. Even at this lower dose of neratinib (figure 1D), we again found no statistically significant improvement for the neratinib plus fulvestrant combination versus single agent neratinib (p=.73). We acknowledge that both of these PDX’s had 3-5 prior lines of hormonal therapy and the patient from whom WHIM51 was established had been previously treated with fulvestrant. Immunohistochemistry on WHIM51 showed that only a few percent of the xenograft cancer cells are ER positive (figure 2A), which may also contribute to the limited effect of single agent fulvestrant on these samples.

**Figure 1.**
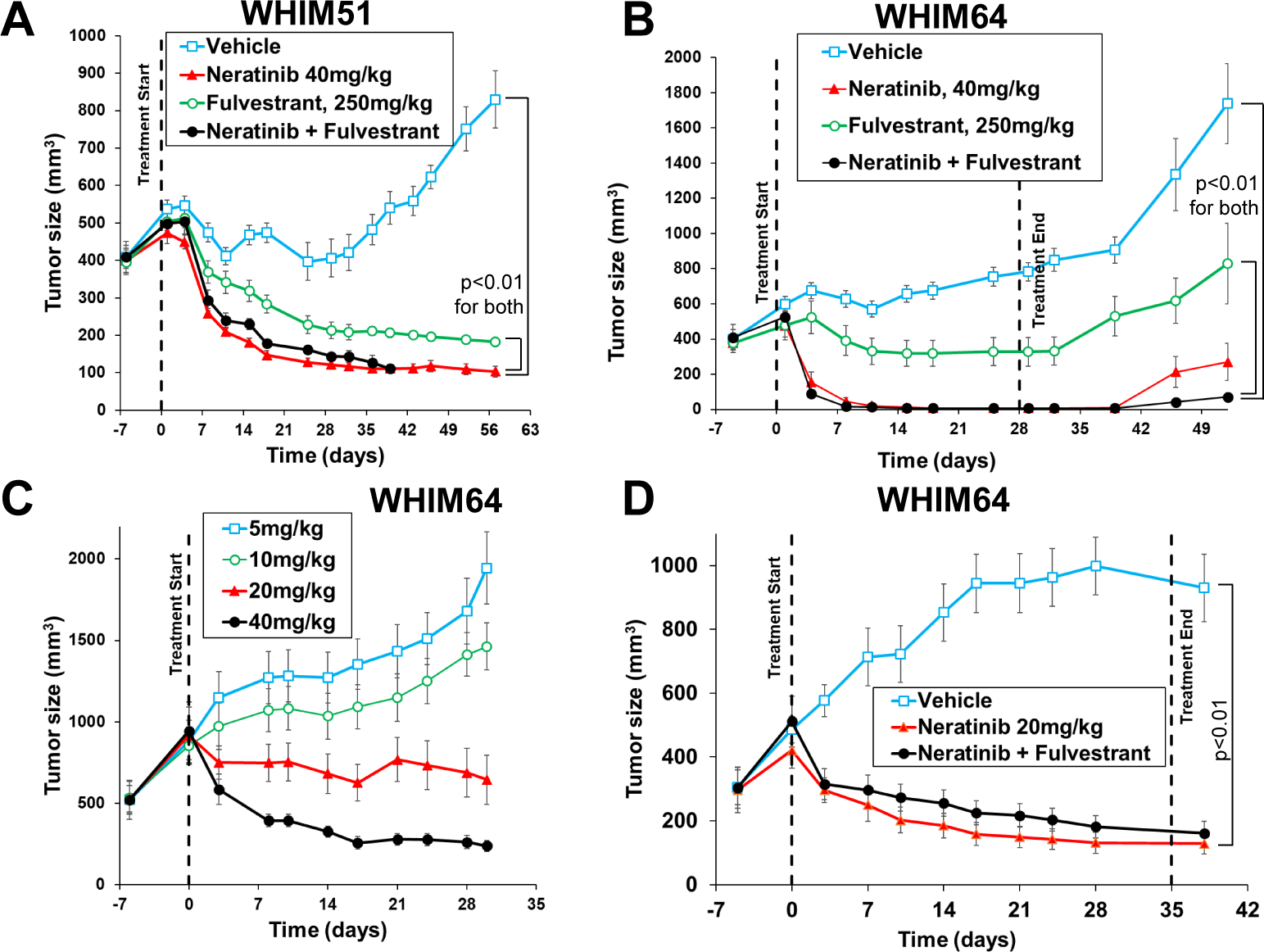
HER2 mutated breast cancer PDX’s. **A,** Effect of neratinib 40mg/kg and/or fulvestrant on WHIM51. **B,** Effect of neratinib 40mg/kg and/or fulvestrant on WHIM64. **C,** Neratinib dosing on WHIM64. The 20mg/kg neratinib dose was chosen for the subsequent drug combination experiments. **D,** Neratinib 20mg/kg alone or in combination with fulvestrant in WHIM64. For A-D, data is mean ± SEM for n=5 mice per arm.

**Figure 2.**
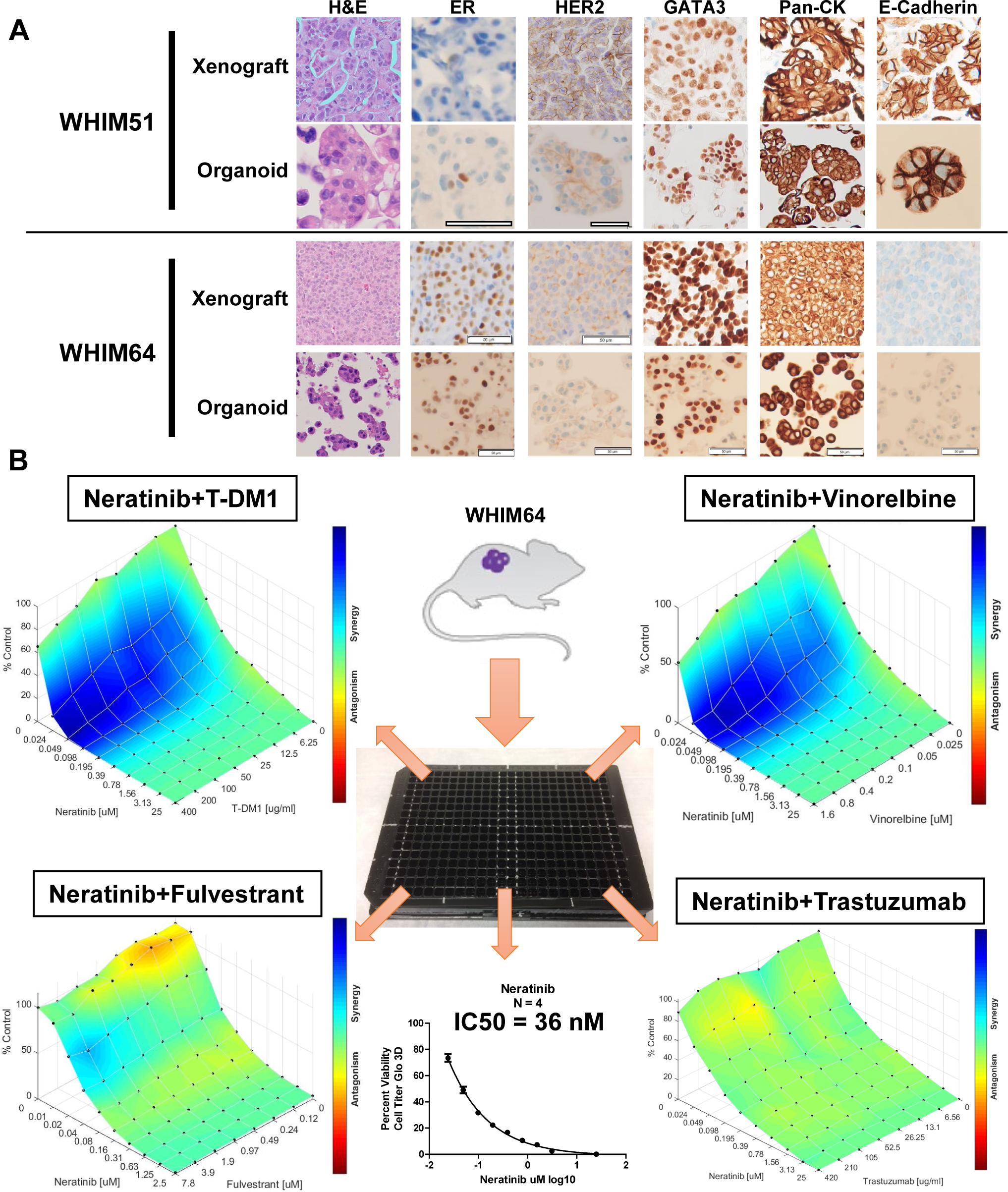
Immunohistochemistry and *ex vivo d*rug combination testing on organoids. **A,** Comparison of histology between PDX tumors *in vivo* and their organoids grown *ex vivo*. Pan-CK = pan-cytokeratin. Scale Bars = 50 microns. **B,** Four drug doublets were performed on a single 384 well plate in a 6 day cell viability assay. Vertical axis represents cell viability as measured by Cell Titer Glo. Three independent replicates of this experiment were performed.

### Organoids from Patient-Derived Xenografts

These results with neratinib plus fulvestrant indicated to us that more drug combinations need to be tested for HER2 mutated MBC. However, these ER+ breast cancer PDX’s are slow-growing and require 4 to 6 months for each drug dosing experiment, therefore, in order to accelerate progress, we tested the ability of these PDX’s to grow as organoids *ex vivo*. Clevers and colleagues published the methodology to grow breast cancer organoids (17) and we found that both of these HER2 mutated PDX’s grew well in breast cancer organoid culture. To prove that these cultures are human breast carcinoma cells, we used immunohistochemistry and PCR markers. Immunohistochemical staining for the mammary epithelial transcription factor, GATA3, and the epithelial cell marker, pan-cytokeratin, was performed (Fig. 2A). Both organoids from WHIM51 and WHIM64 are positive for these two marker proteins, matching the xenografts grown *in vivo*, and indicating that these are mammary epithelial cells. PCR was used to verify the cells are of human origin rather than being murine (supplementary figure S2A). The E-cadherin immunohistochemistry was performed. Organoids from WHIM51, which is an invasive ductal breast carcinoma, express strong membranous E-cadherin, whereas organoids from WHIM64, which is an invasive lobular breast carcinoma, do not. The findings match the E-cadherin expression in the corresponding xenograft tissue sections, indicating the organoids retain the specific ductal or lobular differentiation (Fig. 2A).

### Drug Testing on HER2 Mutated, Breast Cancer Organoids

These organoids were grown for up to 5 passages in this *ex vivo* culture. Drug testing was done on passage zero or passage one organoids in order to minimize the possibility of genetic drift of the cells in culture, avoid overgrowth by normal mouse cells, and to increase the speed of experiments. In order to reduce the number of possible drug combinations to test, we chose to focus on neratinib combinations that had human safety data from phase 1 or 2 clinical trials. Neratinib plus ado-trastuzumab emtansine (T-DM1) and neratinib plus trastuzumab combinations have been recently investigated in clinical trials with patients with HER2+ breast cancer (24,25), a phase 1/2 trial of neratinib plus vinorelbine was published by Awada et al (26), and neratinib plus fulvestrant was tested in HER2 mutant breast cancer patients in the MutHER and SUMMIT trials (9,10).

We dissociated a WHIM64 tumor and plated the cells in a 384 well plate (figure 2B). This plate was configured so that each quadrant of the plate gave an 8 x 10 matrix of the indicated two drugs. Drug synergy was calculated using the Loewe model implemented in the Combenefit software (27). This software graphed the drug combination results as a dose-response surface plot, where the neratinib dose is indicated on the X-axis, the dose of the second drug is indicated on the Y-axis, and the cell viability, as measured by Cell Titer Glo, is indicated on the Z-axis. This surface is colored with a heat map indicating drug synergy in blue, additivity in green, and drug antagonism in red.

We found strong drug synergy with both the neratinib + T-DM1 and neratinib + vinorelbine combinations, as indicated by the large, blue colored regions on the drug combination dose-response surface plots (fig. 2B). In contrast, the neratinib + fulvestrant and neratinib + trastuzumab combinations showed primarily additivity and did not have comparable, large regions of drug synergy on these plots. The limited effect of fulvestrant seen here matches the *in vivo* result shown in figure 1C. The patient from whom WHIM64 was derived had three prior lines of hormonal therapy and therefore, WHIM64 is likely to be a hormone refractory breast cancer. The effect of trastuzumab could be underrepresented in this breast organoid culture, because no immune effectors are present in this system and immune-mediated mechanisms of action of trastuzumab are an important component of its efficacy (2).

We performed similar drug treatment experiments on WHIM51 organoids (Figure 3). WHIM51 forms solid clusters as organoids and is harder to plate uniformly in 384 well plates, therefore, we performed WHIM51 organoid experiments in 96 well plates. Extending the range of drug beyond an 8 x 10 matrix was done by using multiple 96 well plates to cover the larger range and including vehicle controls and blanks on every plate. Like WHIM64, WHIM51 organoids showed drug synergy when treated with neratinib + T-DM1 and neratinib + vinorelbine, as indicated by the large blue areas in Figures 3A, B. No synergy was seen with the neratinib + fulvestrant or neratinib + trastuzumab combinations (Fig 3C, D). As noted before, WHIM51 has weak expression of ER and the patient from which it was obtained had prior exposure to fulvestrant, as well as multiple other hormonal therapy drugs (figures 2A and 1A), explaining the poor activity of fulvestrant on this sample. Similarly, lack of immune components in organoid culture conditions could explain the lack of activity of trastuzumab seen here.

**Figure 3.**
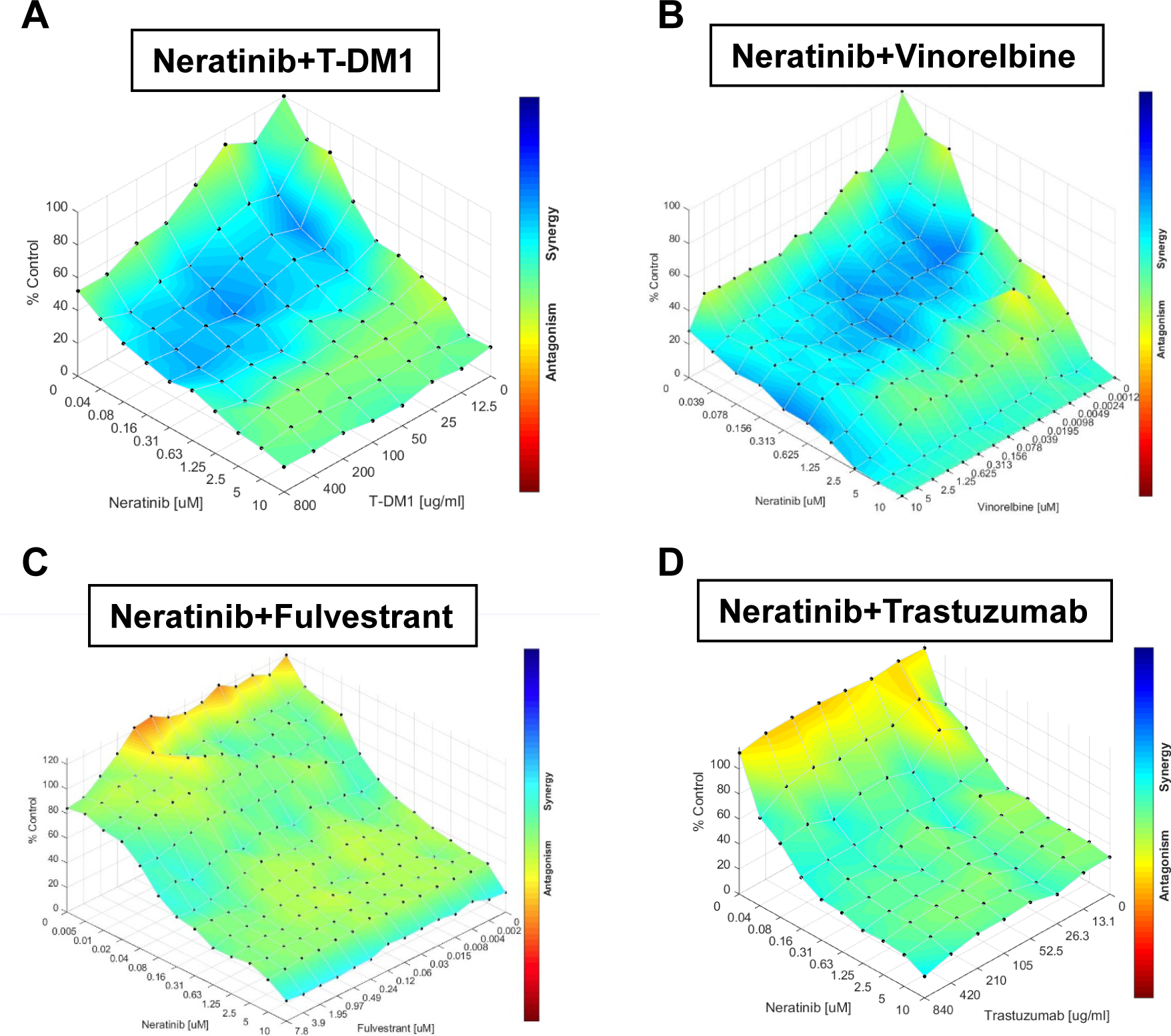
*Ex vivo* drug combination testing on WHIM51 organoids. Combination dose-response surface plots are as per figure 2. Three independent replicates of all drug combinations were performed.

### *In vivo* Confirmation and Mechanism of Drug Synergy

WHIM64 and WHIM51 were implanted into NSG mice to confirm these *ex vivo* organoid culture results (Figure 4). Single agent T-DM1 slowed the rate of WHIM64 tumor growth and stabilized the tumor size at about 750 mm^3^, single agent neratinib (20mg/kg) had a static effect on tumor size, and their combination produced statistically significant tumor regression (Fig. 4A). In WHIM51 tumor (Fig. 4B), neratinib 20mg/kg daily prevented tumor growth for 21 days, but then rapid tumor growth was seen. T-DM1 monotherapy caused tumor regression, but tumor regrowth occurred after day 42 despite ongoing weekly dosing. The combination showed tumor regression that was sustained through the end of the 2 month experiment (Fig 4B). Similarly, neratinib + vinorelbine treatment caused tumor regression of WHIM64 and WHIM51 (Fig. 4C, E), which was a stronger effect than either drug on its own. Vinorelbine monotherapy slowed the growth of WHIM64 (Fig 4D), whereas it had a stronger effect on WHIM51 (Fig 4E). In sum, strong tumor regression was seen for both the neratinib + T-DM1 and neratinib + vinorelbine combinations, matching what was seen in the organoid culture experiments (Fig. 2B, 3).

**Figure 4.**
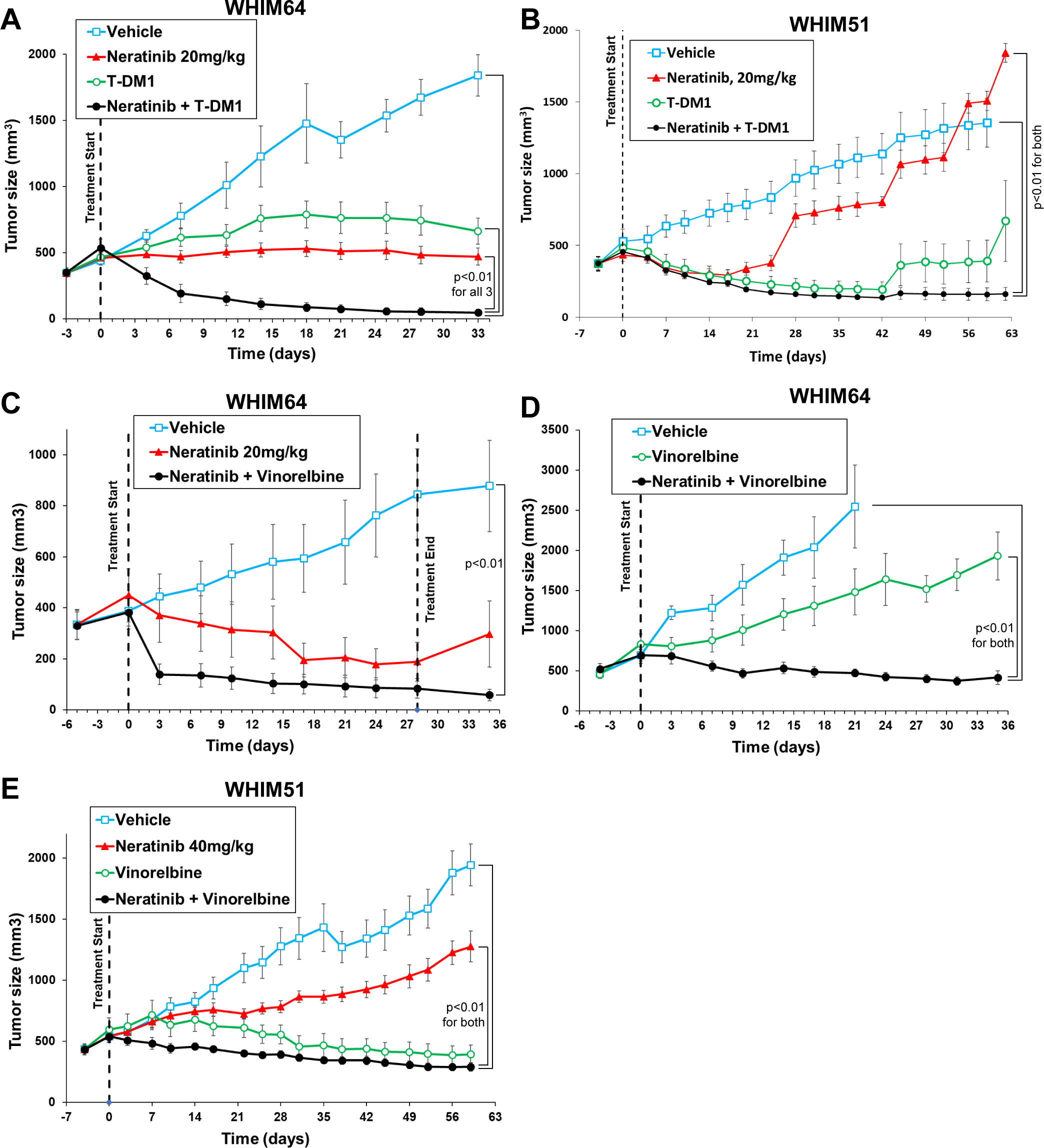
*In vivo* testing of drug combinations. **A-B,** Effect of neratinib 20mg/kg po daily and/or 10mg/kg T-DM1 IV weekly on WHIM64 (A) and WHIM51 (B). **C-D,** Effect of neratinib 20mg/kg po daily and/or vinorelbine 10mg/kg IV weekly on WHIM64. **E,** Effect of neratinib 40mg/kg po daily and/or vinorelbine 10mg/kg IV weekly on WHIM51. For **A-E**, data is mean ± SEM for n=5 mice per arm.

Prior literature evidence provides substantial information about the potential mechanisms of drug synergy. Vinorelbine is an anti-microtubule agent that has been combined with HER2 targeted drugs in numerous clinical trials (26,28–30). Anti-microtubule agents, such as taxanes, vinorelbine, and eribulin, are frequently combined with HER2 targeted drugs and trastuzumab has been shown to increase taxane-induced apoptosis in HER2 amplified cell lines by downregulating the Cdc2 inhibitor protein, p21^Cip1^ (31,32). As a control, we tested neratinib with doxorubicin on WHIM64 organoids, as WHIM64 is sensitive to doxorubicin and doxorubicin does not target microtubules. We did not observe drug synergy with this combination (supplementary figure S3).

For the neratinib + T-DM1 combination, we investigated how neratinib affected the uptake and internalization of T-DM1 and trastuzumab because neratinib and the closely related, irreversible HER2/EGFR inhibitor, CI-1033, have been shown to increase HER2 internalization via the endocytic pathway (33–35). We used a real-time, live cell fluorescence assay that uses an anti-human Fab labeled with a pH-sensitive fluorophore that has a large increase in fluorescence when localized to endosomal–lysosomal compartments (36). We observed that neratinib treatment increased the uptake of both T-DM1 and trastuzumab in BT474 cells (Fig. 5A – B). This effect does not occur with the reversible HER2 inhibitors, lapatinib and tucatinib (Fig. 5A), but does occur with other irreversible HER2 inhibitors, afatinib and dacomitinib (Fig. 5B). To further study the mechanism of synergism, we treated WHIM64-bearing mice for 1, 3, or 10 days with T-DM1 alone or in combination with neratinib. We found that apoptosis was greatest at day 3 in WHIM64 bearing mice receiving the T-DM1+neratinib combined therapy, as evidenced by the pyknotic cells and apoptotic bodies seen on the H&E stained sections and the cleaved PARP immunohistochemistry (Fig. 5C). At day 10, T-DM1 monotherapy caused only very focal necrosis, whereas the T-DM1+neratinib combined therapy caused extensive cell death (indicated by the open arrow, Fig. 5D). These results demonstrate that neratinib-mediated increased uptake of T-DM1 generates increased tumor cell kill, likely through greater intracellular levels of the DM1 payload to the breast cancer cells.

**Figure 5.**
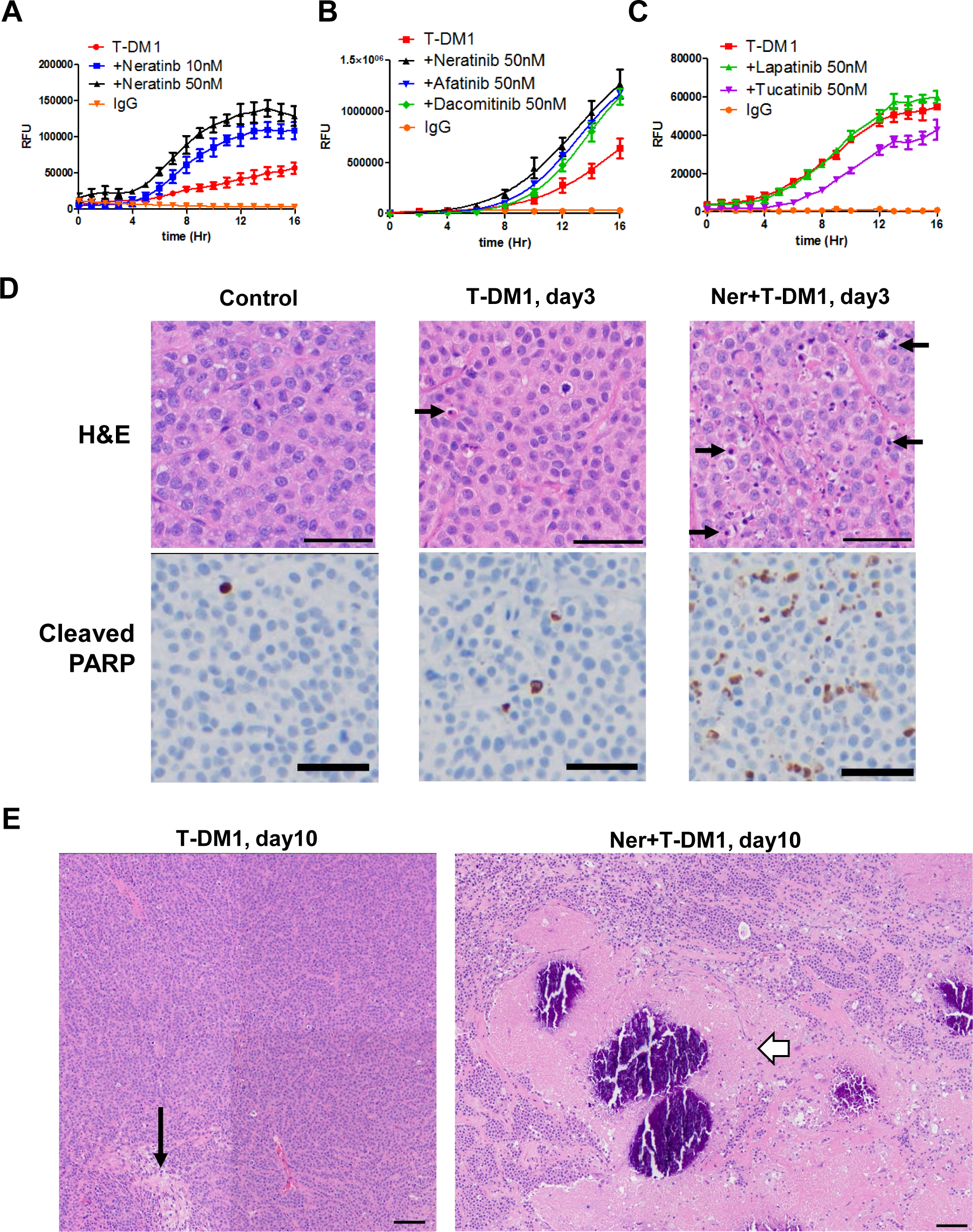
Mechanism of synergy of neratinib plus T-DM1. **A-C,** Uptake of T-DM1 by BT-474 cells as measured using FabFluor Red label. BT474 cells were incubated with control IgG, T-DM1, or T-DM1 plus the indicated tyrosine kinase inhibitors. Real time measurement of intracellular red fluorescence were obtained using a Sartorius Incucyte Zoom. **D**, H&E and Cleaved PARP IHC of WHIM64 treated *in vivo* with the indicated drugs for 3 days. Black arrows indicate pyknotic cells and apoptotic bodies. Scale bar is 50 um. **E,** H&E images of WHIM64 after 10 days of *in vivo* drug treatment. Scale bar is 100 um and images are composite of 4 or 6 overlapping fields of view. Long black arrow indicates area of focal necrosis. Open white arrow shows area of extensive cell death with large eosinophilic areas with central calcification (dark purple).

## Discussion

Organoid culture greatly accelerates drug testing on PDX’s and this work builds on a prior study by Bruna et al. which used short-term culture of tumor cells from PDX’s (18). Testing four drug combinations was performed in under two weeks using organoid culture and PDX material from one mouse was sufficient to generate multiple plates of cells. In contrast, because these ER+ breast cancer PDX’s are slow growing, 4 to 6 months and 30-50 mice are needed to test one drug combination *in vivo*. While we focused on drug combinations with existing human safety data, it would be conceptually straightforward to extend this organoid drug testing to a library screen or to select additional drugs for testing based on genome sequencing or other molecular characterization of the PDX’s. Further, this study gives an example of a tractable way to study cancer drug combinations. Multiple drug combinations can be tested rapidly in organoid culture and we avoided the problem of combinatorial explosion of possible drug combinations (37) by choosing a clinically pragmatic strategy which emphasizes drug combinations that can be rapidly translated into phase II and phase III clinical trials.

These experiments identified two synergistic drug combinations for HER2 mutated, non-amplified metastatic breast cancer. Neratinib plus T-DM1 (ado-trastuzumab emtansine) was synergistic in both WHIM64 and WHIM51 and human safety data on this combination has been published (24). Similarly, the neratinib plus vinorelbine combination is synergistic in both of these PDX’s and human safety data on this combination was published in 2013 (26). HER2 targeted therapeutics are frequently combined with anti-microtubule agents and vinorelbine is an inexpensive and well tolerated chemotherapy agent for metastatic breast cancer patients (28–30). Vinorelbine has advantages over taxanes for HER2+ mutated MBC patients in that most of these patients will have previously received a taxane, either in the adjuvant or metastatic setting, and vinorelbine is effective in the third-line or higher setting for metastatic breast cancer (38,39). Further, oral vinorelbine is approved outside of the U.S. (40) and therefore, a combination of two oral agents is a potentially convenient approach for drug therapy.

We acknowledge that there are limitations of this study. First, this study uses only two independent patient samples, but HER2 mutated breast cancer is a rare molecular subtype and efforts to establish or obtain other HER2 mutated breast cancer PDX’s have not yet been successful. Testing the neratinib + T-DM1 and neratinib + vinorelbine regimens on additional HER2 mutated breast cancer PDX’s should be performed once those samples become available. Second, organoid culture does not contain components of the immune system and therefore, these experiments could underestimate the efficacy of trastuzumab. Adding T-cells to organoid culture has been investigated by other groups and could be important for future studies (41,42). Third, organoid culture lacks other cell types that are present in the tumor microenvironment. Addition of cancer-associated fibroblasts to pancreatic cancer organoids has been studied by Tuveson and colleagues (43,44).

Despite these limitations, we feel this work represents a significant advance. Worldwide, over 500 breast cancer PDX’s exist (19) and when considering all cancer types, thousands of PDX’s are available (20,21). Performing high-throughput drug screening on PDX derived organoids (PDxO’s) offers a way to dramatically accelerating drug testing on these patient-derived cancer samples. Drug combinations could be evaluated rapidly, at large scale and in a format that has low input requirements for both tumor material and drug quantities.

In conclusion, organoid culture was used to rapidly test drug combinations across two HER2 mutated MBC patient derived samples. Neratinib + T-DM1 and neratinib + vinorelbine were found to be synergistic combinations and these results were validated *in vivo*. Further, while cancer organoids made directly from patient samples offer great promise for the future, another ready source of human cancer organoids are existing PDX’s. Given the thousands of PDX’s already available worldwide, PDX derived organoids (PDxO’s) offers a way to dramatically accelerating cancer drug testing.

## Methods

### PDX Methods and Drug Dosing

Human tumor samples were collected for implantation into NSG mice (The Jackson Laboratory, Stock No. 005557) and establishment of PDX’s under an IRB approved protocol (Washington University Human Research Protection Office protocol # 201102244). PDX’s were grown as per our prior publication (45) under a Washington University IACUC approved protocol. Drug administration to PDX bearing mice was performed as follows: Neratinib (from Puma Biotechnology) was dosed daily by oral gavage using 0.5% methylcellulose + 0.4% Tween-80 vehicle. Fulvestrant, vinorelbine, and T-DM1 were obtained from the Siteman Cancer Center pharmacy. Fulvestrant was dosed at 250mg/kg twice weekly IM. T-DM1 and vinorelbine were dosed at 10mg/kg by weekly IV (tail vein) injection. Vehicle groups were dosed with 0.5% methylcellulose + 0.4% Tween-80 daily by oral gavage.

### PDX Organoid Culture

Human breast cancer PDX tumors were harvested at 1 to 2 cm^3^ and minced finely (∼1-2mm^3^) to be processed for ongoing organoid culture and assays, or for cryopreservation for future use in 90% FBS/10% DMSO/ 10uM Y-27632. The minced tumors were digested with 5ml to 10ml of 1 mg/ml Collagenase 3 (Worthington cat. no. LS004183) plus 0.5mg/ml Hyaluronidase (Sigma-Aldrich H3884) and 10uM Y-27632 (Selleckchem) in Ad-DMEM/F12+++ (Advanced DMEM/F12 (Life Technologies, cat. no. 12634-010), supplemented with 1x GlutaMAX (Life Technologies cat no. 35050-061) Penicillin-streptomycin (Life Technologies, cat. no. 15140-122), 10 mM HEPES (Life Technologies, cat. no. 15630-080), filtered sterilized (Millipore Steriflip 0.22micron). The tissue was digested overnight at 37°C on a rotating platform. The long digestion is appropriate for breast cancer PDX tumors, but shorter digestions of 1-4 hours are preferred for breast cancer surgical samples. Following digestion, 25 ml of DMEM/10% FBS was added to the mixture as a first wash and the digested tissue was centrifuged at 200g for 5 minutes. This was followed by two more washes of the pellet using 25-30 ml of serum-free DMEM, and the final wash with 25-30 ml of Ad-DMEM/F12+++. If necessary, contaminating RBC were removed by treatment with filtered lysis buffer according to the manufacturer’s instructions (Invitrogen eBioscience™ 1X RBC Lysis Buffer cat no. 00-4333-57). The processed tumor material was either cryopreserved in 90% FBS/10% DMSO/ 10uM Y-27632 for future culture or assay use, or put directly into culture or assays. For organoid culture, the processed tumor tissue was suspended in 50% to 70% growth factor reduced Matrigel (Corning ref 35623) in chilled breast organoid media, as per (17) modified with 10% Noggin/Rpondin1 conditioned media and 5uM Y-27632. The Noggin/Rspondin1 lentiviral expression vector was a kind gift of Blair Madison and Anil Rustgi (46).

The Matrigel suspension of tumor material was deposited onto pre-warmed ultra-low adherence 6-well plates (Greiner cat. no. 657970) in ∼120ul/well distributed as multiple 7ul droplets and allowed to gel at 37°C for 30 minutes and then fed 2ml of organoid media per well. Organoid cultures were also grown in 24-well ultra-low adherence plates (Greiner cat. no 662970) as a single 30ul droplet/well and with 500ul organoid media. Cultures were fed 2-3 times per week and passaged every 7 to 10 days.

### 96-well and 384-well drug IC50 and drug combination assays

Drug IC50 and combination assays were set up using both organoids in culture and tumor material freshly isolated from the PDX or cryopreserved PDX material. Organoids in culture were isolated from Matrigel by treatment at 4°C with Corning Cell Recovery Solution (cat. no. 354253), according to the manufacturer’s instructions. Solid organoids, such as WHIM51, needed treatment with 500ul TrypLE (Life Technologies cat.no. 12604-013) to break the organoids into smaller structures before using them to seed assay plates. The TrypLE digestion along with triteration with a p1000 pipet, was monitored closely to prevent over digestion since WHIM51 will die if processed to single cells. TrypLE was removed by two 12ml washes in Ad-DMEM/F12+++. WHIM64 is a loosely aggregated organoid that disperses with pipetting using a 10ml pipet holding a p1000 tip. Both WHIM51 and WHIM64 were passed through a 100um cell strainer to remove very large organoids before plating for assays. For the 96-well plate (Corning cat.no.3903) assays, the tumor material was centrifuged (200g, 5 minutes) and the pellet suspended in chilled organoid media plus 60% Matrigel. An electronic pipettor (Matrix 250ul) was used to mix and dispense 10ul per well in a ring formation (47). The Matrigel was allowed to gel at 37°C for 30 minutes, then the cultures were fed 100ul/well of breast organoid media. The 96-well plate organoid cultures were incubated 2-4 days before the addition of drug(s) for IC50 or combination assays. Drugs were serially diluted in DMSO, if aqueous insoluble (Neratinib, Fulvestrant), before the addition to organoid media, or serially diluted in media directly, if solubility was not an issue (trastuzumab, ado-trastuzumab, vinorelbine). Intermediate drug concentrations in organoid media were prepared at 2X (IC50 assay) or 4X (drug combination assay). For 384-well plate (Greiner cat. no. 781091) IC50 assays (48), drug in organoid media was prepared at 2X and 30ul was added per well using an Integra Voyager II 2-125ul, 12-channel pipettor capable of changing spacing from 96-well to 384-well plate formats. The tumor material was suspended in chilled organoid media plus 6% Matrigel and 30ul was added per well. For 384 well combination assays, 20ul of each drug was added to 20ul of tumor material in Matrigel. Tumor material was suspended in chilled organoid media plus 10% Matrigel, for a final Matrigel concentration of 3.3%. The drug combination assays in both 96-well and 384-well used a grid format (49) for testing the two drugs. Organoid cultures for IC50 and combination assays were exposed to drug(s) for 6 days. Cell viability was measured with Cell Titer Glo 3D (Promega cat.no G9683) and luminescence was measured using a Tecan infinite M200 plate reader. IC50 values were calculated using the 4-parameter, non-linear regression function in Graphpad Prism 5.0 for Windows. Drug synergy in the combination assays was calculated using the freeware program, Combenefit version 2.021 (27).

### PCR Assay for Human vs Mouse cells

Mouse PIK3CA gene was detected with the following primers: CTT CCC CTT CGC CAA GTC TAA GC (forward) and CAG TTT GAG CTG AAA CCC TGC (reverse). Human PIK3CA gene was detected with CTG TGA ATC CAG AGG GGA AA (forward) and TGG GTA GAA TTT CGG GGA TA (reverse).

### Immunohistochemistry

Organoids were isolated from Matrigel using Cell Recovery Solution, and fixed in 10% neutral buffered formalin for 24 hours. Organoids were then embedded in melted Histogel (ThermoScientific cat.no. HG-4000-012), submitted to the Washington University Histology Core for embedding in paraffin and sectioning. Immunohistochemistry was performed using standard protocols on the Ventana Automated Stainer (Ventana Medical Systems, Tucson, AZ, USA) using Biotin-free multimer technology. Antigen retrieval was done at 95°C with CC1 buffer (Cell Conditioning 1; EDTA-based buffer pH 8.0, Ventana) for 16 to 40 minutes, depending on antibody. The following antibodies were all purchased from Ventana: anti-ER (rabbit monoclonal, clone SP1); anti-Her2 (rabbit monoclonal, clone 4B5); anti-E-cadherin (rabbit monoclonal, clone EP700Y); anti-GATA3 (mouse monoclonal, clone L50-823); and anti-pan cytokeratin (mouse, clones AE1/AE3/PCK26).

### Antibody uptake in BT474 cells

A black wall, clear bottom 96-well tissue culture plate (Falcon cat.no 353948) was seeded with 25,000 BT474 cells per well in 100ul of RPMI with 10% FBS and pen/strep. BT474 cells were purchased from ATCC and authenticated by STR profiling. Trastuzumab and T-DM1 were each diluted to 1mg/ml in PBS. Human IgG, 1mg/ml (BioLegend cat no. 403501) was selected as a negative control for antibody uptake. These antibodies were labeled with the FabFluor Red Antibody Labeling Reagent, as described in (36). One day after seeding, the BT474 cells were treated with Neratinib or DMSO vehicle diluted in media plus FabFluor Red labeled antibodies. Antibody uptake was visualized by the increase of red fluorescence within the cells over 2 days, measured with a Sartorius Incucyte Zoom (Essen BioScience, Ann Arbor, MI).

### Statistical Analysis of PDX Experiments

Tumor volume (TV) was log transformed for better conformity to normality. The linear mixed effects model for repeated measures model was fit on the logged TV data (starting from day of treatment, day1) with the effects of treatment, time (as a categorical variable), and the treatmet*time interaction to estimate the overall treatment effect difference across time and the end-of-study treatment effect at the last available time point based on the least square mean difference and its associated standard error, raw P value and Tukey adjusted p-values. The studentized residual plots from linear mixed effects model were examined for models’ goodness-of-fit.

## Acknowledgements

We thank Drs. Sylvia Boj, Bryan Welm, Katrin P. Guillen, and Jennifer Rosenbluth for technical guidance and helpful discussions on organoid methods. We also thank Dr. Robert Pufahl for PCR assay methods on human versus murine PIK3CA gene and McKenna Wilhelm for performing 384 well experiments on WHIM64.

