## Supplemental Figures and Table for "Neratinib Synergizes with Trastuzumab Antibody Drug Conjugate or with Vinorelbine to Treat HER2 Mutated Breast Cancer Patient Derived Xenografts and Organoids"

**Supplementary Table S1.** Vinorelbine Dosing of NSG strain mice. Vinorelbine was administered IV by tail vein injection weekly x 2 in order to determine the maximally tolerated dose (MTD). Vinorelbine MTD was 10mg/kg for NSG strain mice.

| Vinorelbine dose | Result |
|---|---|
| 1.2 mg/kg | 0/4 mice died |
| 2.4 mg/kg | 0/4 mice died |
| 4.8 mg/kg | 0/4 mice died |
| 10 mg/kg | 0/4 mice died |
| 15 mg/kg | 2/5 mice died and 1/5 mice was hunched and looked sick after the second dose |

Supplementary Figure S1.

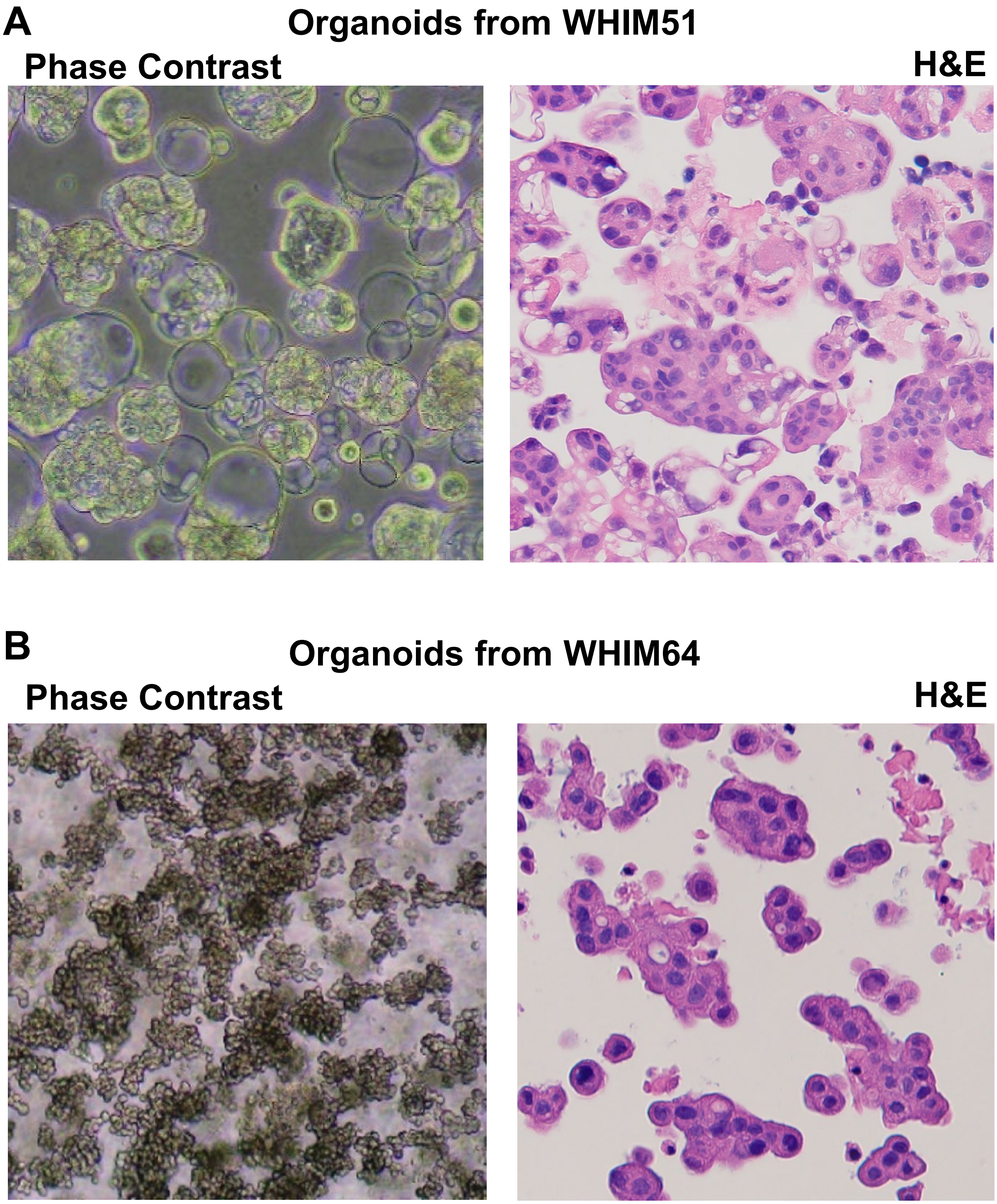


**Supplementary Figure S1. Phase contrast and H&E images of Organoids. A,** WHIM51 forms organoids with a mix of solid and cystic clusters. **B,** WHIM64 forms organoids with cells in grape-like clusters.

**Supplementary Figure S2.**

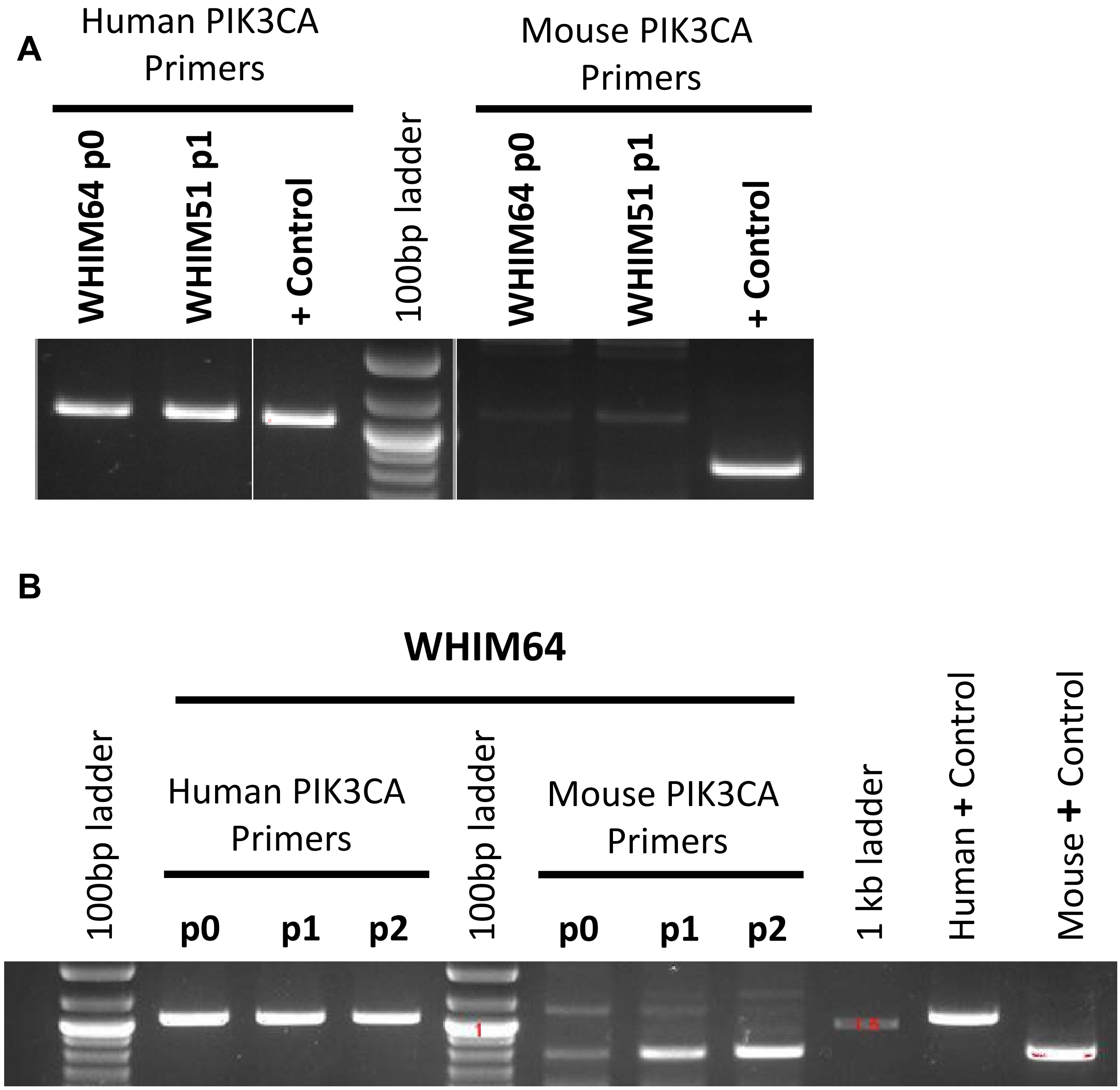


**Supplementary Figure S2. Detection of human and murine DNA in PDX organoid cultures by PCR. A,** PCR to human or mouse PIK3CA gene was performed. Expected sizes of PCR products: 1,085 bp for human and 831 bp for mouse. 100bp DNA ladder shows a heavier band at 1,000 bp. Mouse DNA was very low in passage 0 WHIM64 and passage 1 WHIM51 organoid cultures. B, Same as A. Increasing amount of mouse DNA, representing growth of murine cells, is seen in passage 1 and 2 of WHIM64.

**Supplementary Figure S3.**

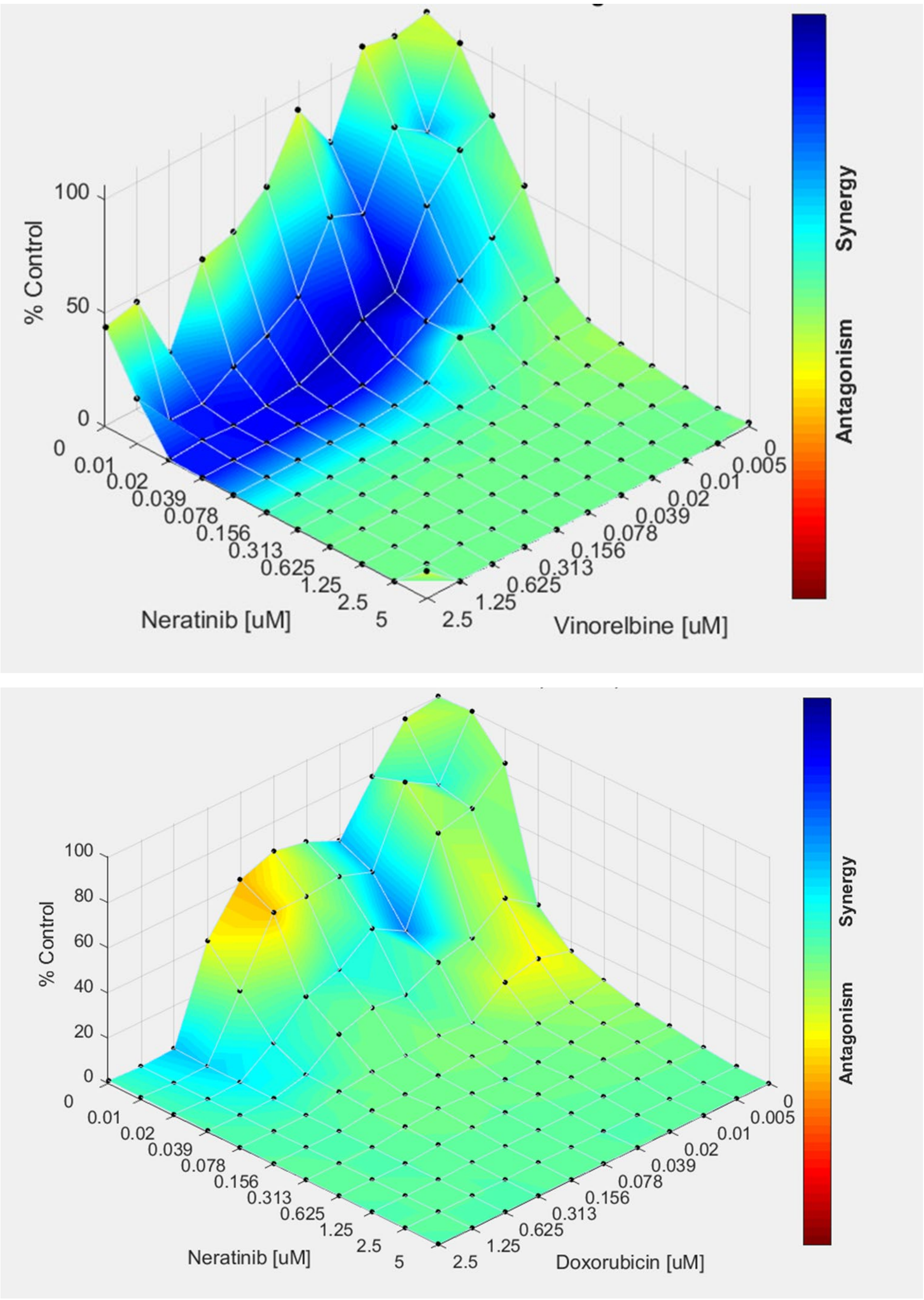


**Supplementary Figure S3. Drug Synergy is seen with Neratinib plus Vinorelbine, but not with Neratinib plus Doxorubicin.** WHIM64 organoids were tested simultaneously in a single 384 well plate in a 6 day cell viability assay. Vertical axis represents cell viability as measured by Cell Titer Glo.

**Supplementary Figure S4.**

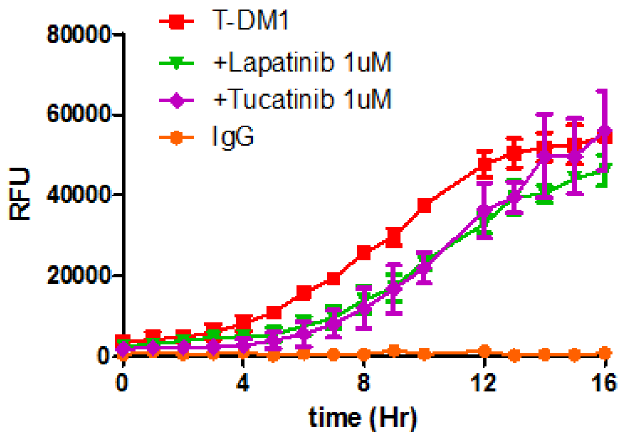


**Supplementary Figure S4.** Uptake of T-DM1 by BT-474 cells as measured using FabFluor Red label and real time fluorescence measurement obtained using a Sartorius Incucyte Zoom. Neither lapatinib 1uM (green triangles) nor tucatinib 1 uM (purple diamonds) increase T-DM1 uptake (compare to T-DM1 alone, red squares).
